# Enhanced semantic classification of microbiome sample origins using Large Language Models (LLMs)

**DOI:** 10.1101/2025.04.24.650461

**Authors:** Daniela Gaio, Janko Tackmann, Eugenio Perez-Molphe-Montoya, Nicolas Näpflin, David Patsch, Lukas Malfertheiner, Christian von Mering

## Abstract

Over the past decade, central sequence repositories have expanded significantly in size. This vast accumulation of data holds value and enables further studies, provided that the data entries are well annotated. However, the submitter-provided metadata of sequencing records can be of heterogeneous quality, presenting significant challenges for re-use. Here, we test to what extent large language models (LLMs) can be used to cost-effectively automate the re-annotation of sequencing records against a simplified classification scheme of broad ecological environments with relevance to microbiome studies, without retraining.

We focused on sequencing samples taken from the environment, for which metadata is important. We employed OpenAI Generative Pretrained Transformer (GPT) models, and assessed scalability, time and cost-effectiveness, as well as performance against a diverse, hand-curated ground-truth benchmark with 1000 examples, that span a wide range of complexity in metadata interpretation. We observed that annotation performance markedly outperforms that of a baseline, manually curated, non-machine-learning keyword-based approach. Changing models (or model parameters) has only minor effects on performance, but prompts need to be carefully designed to match the task.

We applied the optimized pipeline to more than 3.8 million sequencing records from the environment, providing coarse-grained yet standardized sampling site annotations covering the globe. Our work demonstrates the effective use of LLMs to simplify and standardize annotation from complex biological metadata.

## Intro

Streamlining and standardizing metadata is crucial for ensuring reproducibility in scientific research. Over the past decade, the size of the GenBank database has expanded by more than 30-fold, the whole-genome sequencing database (WGS) grew nearly 40-fold, and the European Nucleotide Archive (ENA) reported a 10-fold increase in a window from 2012 to 2022 ^1^. This vast increase underscores the necessity of managing, standardizing, and utilizing such large datasets effectively. Metadata accompanies all scientific data types, and primary data repositories provide submitters with guidelines and facilities for providing structured metadata at the time of submission. However, the submission step–critical as it is–often does not receive as much attention as earlier steps, such as sample collection and processing. Samples with well-organized metadata are more likely to be reused ^2–4^, indicating that thoughtful metadata submission enhances broader research utilization. In the case of raw DNA sequence data, the National Center for Biotechnology Information (NCBI) SRA submission system suggests a number of metadata fields to be filled, along with controlled vocabularies. However, as is common in many databases, only a minimal number of fields are mandatory, leaving submitters with considerable discretion in how information is provided and formatted. This flexibility, while beneficial for submitters, frequently results in metadata that is challenging to reuse. Over time, NCBI has introduced measures to mitigate this issue, such as pre-populated drop-down menus for fields like *organism_type*, and since June 2023, they provide a tutorial to guide submission. While these changes help standardize entries, they are unlikely to completely solve the issue. Moreover, prior submissions that allowed free-form data entry remain difficult to standardize retroactively.

The rise of artificial intelligence (AI) in metadata parsing marks a significant evolution from earlier efforts using traditional Natural Language Processing (NLP) methods ^5–7^ or techniques that use term frequency (*e*.*g*.: term frequency-inverse document frequency, *i*.*e*. TF-IDF) ^8^ to identify key terms within text-based metadata. The complexity of such metadata, which can range from full sentences to acronyms, uses niche terminology and frequently includes spelling variants or typos, posing substantial challenges to traditional NLP and term frequency methods. Relying solely on term frequency, even when metadata is articulated clearly, can be inadequate for good classification, as these methodologies lack contextual understanding ^9^. For example, if both “Komodo dragon” and “mice” are equally mentioned within different, user-defined fields of a metadata text sample, traditional term frequency methods may fail to discern that the first refers to the origin of the sample and the latter to the host’s diet. This limitation highlights the need for more sophisticated approaches. Recent advancements in AI have led to the development of robust pretrained models, particularly Large Language Models (LLMs), which excel at extracting information from diverse and complex metadata across many domains ^10–13^. These models enhance text mining capabilities by effectively understanding context, making them suited for sophisticated tasks like metadata parsing and mining.

We use *MicrobeAtlas* as a testbed for metadata parsing using LLMs. *MicrobeAtlas* is a large, diverse resource, containing millions of metagenomic SRA samples retrieved from NCBI. MicrobeAtlas uses metadata-extracted keywords to assign samples to defined environmental categories based on hard-coded rules. This non-semantic approach can however fail to assign the correct meaning to terms, particularly in the presence of diverse, user-defined metadata fields, leading to ambiguous or even wrong assignments. Our objective here is to instead leverage general-purpose LLMs for (re-)classification of samples into defined environmental categories, while, simultaneously, retrieving other useful information from the metadata. The tasks given to the LLMs are: I. classification of samples into main categories (we here address “biomes” *i*.*e*.: “animal”, “water”, “soil”, “plant”, “other”); II. further classification of samples into sub-categories, here referred to as “sub-biomes”; III. extraction of the geographic location of a given sampling site, and IV. extraction of up to eight key terms describing the sample. We aim to achieve high-quality outputs in a cost- and time-effective manner, exploring the capabilities of GPT under various versions, conditions and configurations. This study addresses a critical gap in working with metadata, by covering a middle ground positioned somewhere between careful, manual re-annotation of a few records on one hand, and using unprocessed metadata of millions of records on the other hand.

## Methods

### Metadata download and processing

We used MicrobeAtlas as a testbed for LLMs metadata extraction. For MicrobeAtlas, the NCBI Sequence Read Archive (SRA) had been searched for DNA sequencing runs with metadata keywords matching “metagenomic”, “microb*”, “bacteria” or “archaea”, and all metadata files for these runs were downloaded. For the present study, all metadata was merged into a single comprehensive metadata file (github/metadata_mining/source_data/sample.info). This file was then organized on a per-sample basis and sorted into directories named after the last three digits of each sample’s name (script: *01_dirs*.*py*). Environmental-, food-, plant-, species-, and cross-species anatomy-ontologies (ENVO, FOODON, NCBITaxon, PO, UBERON) were retrieved, parsed and subsequently consolidated into a dictionary (script: *fetch_and_join_ontologies*.*py)*. Metadata files underwent a cleaning process and ontology terms were converted from their numeric representation to their corresponding textual description, utilizing the ontology dictionary we assembled (script: *clean_and_envo_translate*.*py*). During cleaning, we removed empty lines and fields containing placeholder texts (*e*.*g*.: “missing”, “NaN’, “not applicable”), as well as lines that began with “experiment”, which typically include information about wet lab procedures (*e*.*g*.: sample preparation, sequencing, library preparation, adapter sequences). Log files were maintained in order to track which lines were removed and which ontology codes were converted to their textual counterparts. Coordinates (latitude and longitude) were extracted from the metadata, based on a combination of field names and parseable numeric formats (script: *parse_lat_lon_from_metadata*.*py*). Occurrence frequencies of major biome-indicating terms within NCBI submission fields was computed (script: *field_distrib_analysis*.*py*) and the size of metadata files was monitored to ensure they remained manageable for further processing (script: *check_metadata_sizes*.*py*). (**Figure 1**; **Supplementary Figure 1**)

**Figure 1.**
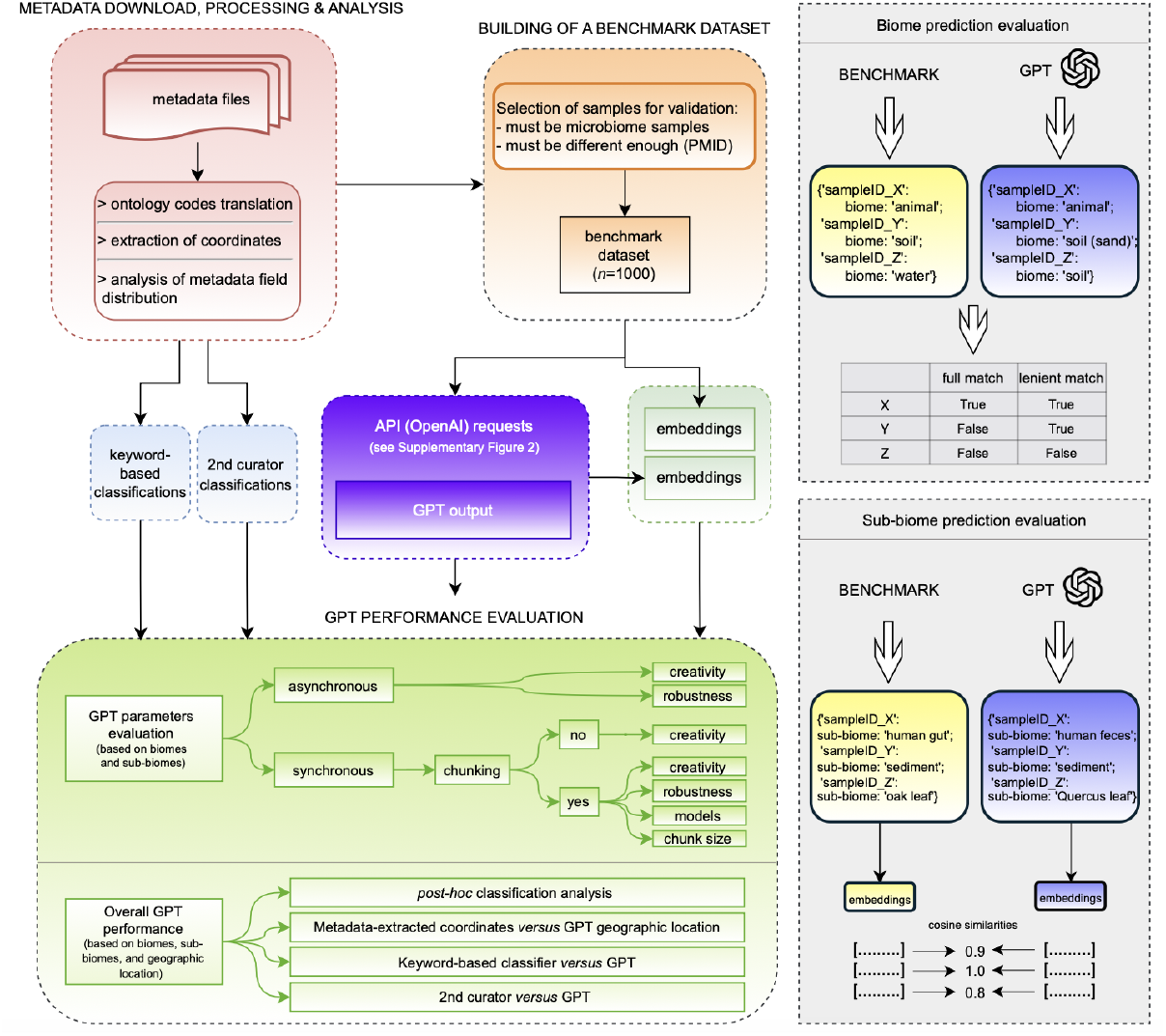
Summarized pipeline. For a detailed pipeline see Supplementary Figure 1.

### Building a benchmark dataset

To assess the output from the LLMs, we established a benchmark dataset consisting of hand-curated metadata records. This process began with a sub-selection of metadata records from the initial pool of 3.8 million biological samples and ultimately selecting 1 million samples based on the availability of sufficient metadata. From these selected samples, we extracted identifiers such as DOI, PMID, PMCID, and BioProject accession numbers. Utilizing the public resource *Entrez*, we retrieved referenced publications for each sample record, retaining all samples for which at least one valid Pubmed abstract could be retrieved. In instances where multiple publications were associated with a single sample, we chose the oldest one, assuming it’s the most likely primary data reference. However, if the title of the publication contained keywords suggesting it described a protocol, we selected the second oldest publication. The primary objective of this publication assignment strategy was to ensure that each sample selected for the benchmark was linked to a unique abstract, thereby maximizing the likelihood that the samples within the benchmark dataset were sufficiently distinct from one another. This approach helps avoid potential biases that might arise if multiple samples from the same study are selected, which would result in redundant metadata. (scripts: *make_subset_large_file*.*py* and *merge_identifiers*.*py*) (**Figure 1**; **Supplementary Figure 1**)

The specific task of the hand curation step was to classify samples into five biomes: ‘animal’, ‘soil’, ‘plant’, ‘water’, and ‘other’, based on all preprocessed metadata. The choice of these biomes was based on *MicrobeAtlas’* pre-existing two-levels ontology, which is what we refer to with “biome” and “sub-biome”. Following *MicrobeAtlas’* choice, to assign “rhizosphere” samples to the plant-, and “sediment” to the water-biome, we did so in the hand curation step, too. Samples with an identifiable origin that did not match any of the five biomes were discarded from the benchmark dataset. Sub-biomes were determined based on more detailed manual examinations of all metadata fields. For animal and plant biomes, sub-biomes correspond to the host taxon and the specific part from which the sample is taken. For example, samples from the animal biome may be classified into the sub-biome “human gut”, while those from the plant biome can be categorized as “olive tree leaf”. Water biome samples were described by their water body source (*e*.*g*.: river, sea, waste water, ocean, lake, etc). Soil samples were differentiated by soil type (*e*.*g*.: agricultural, forest, tundra, desert, peatland, etc). Samples with biome category “other” were sorted into the following most frequently encountered sub-biomes: urban, bioreactor, laboratory, feed/food, fungus, air, or other. The sub-biome was assigned by the curator as it was reported in the metadata. For example, if the sample host was described using its scientific or its common name, it was assigned accordingly. (**Figure 1**; **Supplementary Figure 1**)

We describe briefly the “keyword-based classifier” that MicrobeAtlas is currently based on. Keywords are extracted from various fields and processed (*e*.*g*.: lowercasing, removal of special characters, tokenization). Keywords are matched against environment-specific term sets. For example keywords “leaf, banana, tree, crop” match with “plant”. When keywords of a sample were observed to match with more than one environment-specific term set (*e*.*g*.: “leaf, banana, tree, insect”), the sample gets labeled as “unknown”.

To facilitate a balanced selection of samples across biomes when building the benchmark dataset, we leveraged MicrobeAtlas biome classifications based on the “keyword-based classifier” described above (script: *make_gold_dict*.*py*). This script retrieves sample metadata, enabling the curator to dynamically assign the most appropriate biome and sub-biome. To allow overrides of biomes or sub-biomes, the script *edit_gold_dict*.*py* was employed. Additionally, sub-biomes were used in *field_distribution_analysis*.*py* where they were matched (full and partial matches) against the metadata, in order to establish which fields within the metadata were informative of the sample origin. (**Figure 1**; **Supplementary Figure 1**)

### Requests to GPT

For synchronous requests to GPT, a pipeline was built (script: *openai_main*.*py*), structured into five components: (1) setup of a logging system; (2) fetching metadata texts and segmenting them into manageable chunks; (3) executing requests to the OpenAI API; (4) an initial parsing of the GPT output, during which sample IDs between input and output are matched. Step (4) allows sample IDs missing from the output or that failed parsing to be requested again. (**Supplementary Figure 2A**)

For asynchronous interactions with OpenAI, the pipeline involves retrieving the metadata texts (script: *metadata_processing*.*py*), sending batch requests through the API (script: *gpt_async_batch*.*py*), and retrieving the output up to 24 hours later (script: *gpt_async_fecth_and_save*.*py*). (**Supplementary Figure 2B**)

Both synchronous and asynchronous interactions generate a consolidated output file. The name of the output file reports various parameters, such as the number of metadata samples per biome (`--nspb`) randomly selected from the benchmark dataset, and reproducibility via a fixed random seed (`--rs`). Synchronous runs allow for chunking (`--chunking`), where a single request contains the system prompt followed by multiple metadata samples (if chunking is enabled) or a single sample (if disabled). The maximum number of samples fitting into a single chunk depends on chunk size (`--chunksize`), which reflects the total number of tokens that can fit into a single request, including the system prompt. We employed a first-fit decreasing binning mechanism to optimize chunk utilization. This method sorts samples by token count in descending order and packs them into the fewest number of chunks possible, ensuring that each chunk is filled close to its token capacity without exceeding it (script: *openai_02_metadata_processing*.*py*). Asynchronous requests are never chunked, meaning that one sample metadata per request is sent. Furthermore, users can specify the model (`--model`), the maximum token count (`--maxtokens`), and parameters that determine creativity of chat completion, *i*.*e*.: temperature (`--temp`), nucleus sampling (`--topp`), frequency penalty (`--freqp`), and presence penalty (`--presp`). The output filename also includes optional text (`--opt_text`) optionally detailing specifics of the run, the total number of API requests (for synchronous requests), and the timestamp of the output file creation. The designation “batch” within the filename indicates the OpenAI-generated unique ID of an asynchronous request.

To ensure uniformity across all interactions with OpenAI API, a standardized system prompt was used. This prompt was designed to systematically collect specific information for each sample from its metadata in a structured manner, so as to facilitate a streamlined parsing and analysis of the output. The system prompt was structured as follows:

*We kindly request your expertise in analyzing the following microbial metagenomic samples from their metadata texts:*

- *Please deduce the source category for the sample, choosing from ‘animal’ (including humans), ‘plant’, ‘water’, ‘soil’, or ‘other’. Your choices are: ‘animal’ (incl. human), ‘plant’, ‘water’, ‘soil’, ‘other’. Give strictly a concise, single-word label for the sample ID*.
- *Please infer geographical location where the sample was collected, including the country (NOT the coordinates)*.
- *Extract strictly 5 to 8 keywords descriptive of the sample origin, separated by commas. Put them within curly brackets*.
- *We seek a brief, up to three-word description of the sample’s specific origin. For ‘animal’ or ‘plant’ sources, please specify the host and part thereof. For ‘water’ samples, the type of water body is sought (e*.*g*., *lake, brine, sea, waste water, etc). If from ‘soil’, specify (e*.*g*.: *agricultural, desert, forest, etc). If from ‘other’ specify which (e*.*g*.: *urban, laboratory, feed/food, fungus, air, etc)*.

*If information is missing, kindly indicate ‘NA’. Please separate all values with three underscores (‘ ‘)*.

*An example response: SRS123456 animal Los Angeles, USA* {*medical, bone fracture, infection, collagen, hospital, intensive care, cast, Staphylococcus epidermidis*} *human elbow*

The prompt above was used in all cases except for asynchronous requests and synchronous requests that are used for a direct comparison with asynchronous requests. For these cases, GPT is prompted to generate the output in a json format (see: [github/metadata_parsing/source_data/openai_system_prompt_json.txt](https://github.com/GaioTransposon/metadata_mining/blob/main/source_data/openai_system_prompt_json.txt)).

To improve the accuracy of GPT classifications for challenging cases (*e*.*g*.: rhizosphere and sediment samples), we implemented a tailored prompt (see: [github/metadata_parsing/source_data/openai_system_better_prompt.txt](https://github.com/GaioTransposon/metadata_mining/blob/main/source_data/openai_system_prompt_json.txt)). This prompt instructs GPT to categorize rhizosphere samples as ‘plant’ and sediment samples as ‘water’. In addition, a human curator was engaged to assess sample classifications using the standard prompt initially and the improved prompt in a subsequent round. This approach allowed us to directly compare the performance of GPT against human curation, particularly for these challenging sample types, and to verify whether targeted prompting improves sample classification.

We conducted a series of tests to compare synchronous and asynchronous request methodologies. For synchronous requests, we examined the impact of enabling chunking versus disabling it, and we assessed how varying the chunk sizes affect the output. Additionally, we evaluate different models, robustness, alongside adjustments in creativity parameters of chat completion, which can influence, among others, term repetition. In scenarios where chunking is disabled, we focus on the effects of tweaking creativity parameters. Our rationale is that, while concatenating multiple metadata texts into a single request might affect output due to term repetition penalties, this effect might differ when each metadata text is sent as a separate request. Similarly, for asynchronous requests, tests to assess performance of different models, as well as their robustness and the effect of tweaking creativity parameters are performed. (**Figure 1**)

### Validation statistics

To validate biome classifications, the output files are parsed to extract the “biome” column. For each sample ID, the biome assigned by GPT is compared with the corresponding biome in the benchmark dataset.. We classify matches in two ways: full matches, where the GPT-assigned biome exactly matches the benchmark biome (recorded as “True” or “False”), and lenient matches, where a partial match is accepted (*e*.*g*.: benchmark dataset: *animal*; GPT output: *animal (human)*). Thus, we analyze full matches and combined-full+partial matches distinctly (the latter referred to as “lenient matches”). In the lenient case, such partial matches are also recorded as “True”. We analyse both full matches alone and full+partial (lenient) matches separately. For comparisons involving repeated sample IDs across runs, we use the Mcnemar test, which is appropriate for paired binary outcomes (True/False). For comparisons across different sample sets, we employ the t-test for independent samples. In both scenarios, a Bonferroni correction is applied to adjust for multiple comparisons. (script: *validate_biomes_subbiomes*.*py*) (**Supplementary Figure 1**)

Sub-biome validation is less straight-forward, given the free-form data entries in the benchmark dataset and the flexible assignments by GPT. The “sub-biome” column is extracted from the GPT output and the text-embedding-3-small model is used to generate embeddings for each sample ID from both the GPT sub-biomes and the benchmark dataset sub-biomes (script: *embeddings_from_sb*.*py*). These embeddings are compared to calculate cosine similarity, providing a quantitative measure of the accuracy of GPT’s sub-biome predictions. (script: *validate_biomes_subbiomes*.*py*) Similarly to biome prediction validation, for runs involving different sample IDs, a t-test for independent samples is used. For runs involving the same sample IDs, comparisons are performed using the paired t-test, which is suitable for comparing cosine similarity scores. (**Supplementary Figure 1**)

In order to assess the performance of GPT in accurately determining the location of sample collection, we use latitudes and longitudes extracted from the metadata (script: *parse_lat_lon_from_metadata*.*py*). These are then geo-coded to textual descriptions using the geopy library (script: *coord_to_text*.*py*). String matches between GPT-predicted locations and the geo-coded locations are recorded. For mismatches, latitude and longitude coordinates for the GPT-predicted locations are obtained using the Google Maps API. This allows us to measure the distance between the GPT-predicted coordinates and those extracted from the metadata. Mismatches are categorized based on the distance between these points into the following categories: less than 100 km, 100-500 km, 500-1000 km, 1000-4000 km, and over 4000 km. Additionally, to determine whether mismatches are due to incorrect GPT predictions or errors in the metadata-derived coordinates, 100 randomly selected samples are manually validated. (script: *geo_check*.*py*) (**Supplementary Figure 1**)

We conduct a comprehensive analysis of GPT’s performance by aggregating outputs across various experimental parameters (*i*.*e*.: chunking, (a)synchronicity of requests, and creativity parameters) (script: *overall_analysis*.*py*). The script compiles all GPT output files, merging them into a dataset for comparative analysis against curator-assigned biomes (benchmark dataset). The lenient accuracy is calculated to assess the accuracy of biome predictions both overall and within specific biome categories. The script also compares the GPT-predicted classifications with MicrobeAtlas’s previous biome predictions (based on the “keyword-based classifier”) to determine whether GPT has improved our sample classification. (**Supplementary Figure 1**)

To contextualize the accuracy of GPT-based classification, we conducted a comparative assessment between machine (GPT) and human performance. We randomly selected 250 samples from the benchmark dataset and provided the corresponding metadata to a trained molecular biologist with no prior exposure to this project. The human annotator was asked to assign biomes and sub-biomes to the samples using the same system prompt initially used for GPT (see: “Requests to GPT” section). In a second round, both the human annotator and GPT were presented with an improved version of the prompt. The revised prompt included a clarifying instruction: “Please note that rhizosphere samples should be categorized as ‘plant’ and sediment samples as ‘water’.” This addition was made to assess whether explicit guidance improves classification accuracy for ambiguous cases. We evaluated and compared the performance of both classifiers--GPT and human--under each prompt condition (standard vs improved). This experiment allowed us to understand how well GPT performs relative to an uninitiated but scientifically literate human, and whether targeted prompting can effectively steer, not only human, but also machine classification.

## Results

### GPT overall performance

When testing the classification performance of our LLM-based classifier (GPT 3.5) on our manually curated benchmark dataset (*n*=1000 samples, 200 per biome), using a carefully crafted prompt (see Methods), we found that the model reached an average biome classification accuracy of 80.6% (precision: 81.7%, F1-score: 80.16). This performance was robust to variations in GPT run modes and parameters (n=125; accuracy: 76.0% to 84.1%; precision: 77.1% to 85.8%; F1-score: 76.1% to 84.0%) and was achieved without any example-driven fine-tuning of the model. When examined separately by biome, the accuracy shows variability. ‘Soil’ and ‘animal’ biome samples show the highest accuracy with accuracy rates of 93.6% and 94.3%, respectively. In contrast, ‘plant’, ‘water’, and ‘other’ biome samples exhibit lower accuracy rates (67.2%, 79.3%, and 67.5%, respectively) (**Figure 2**).

**Figure 2.**
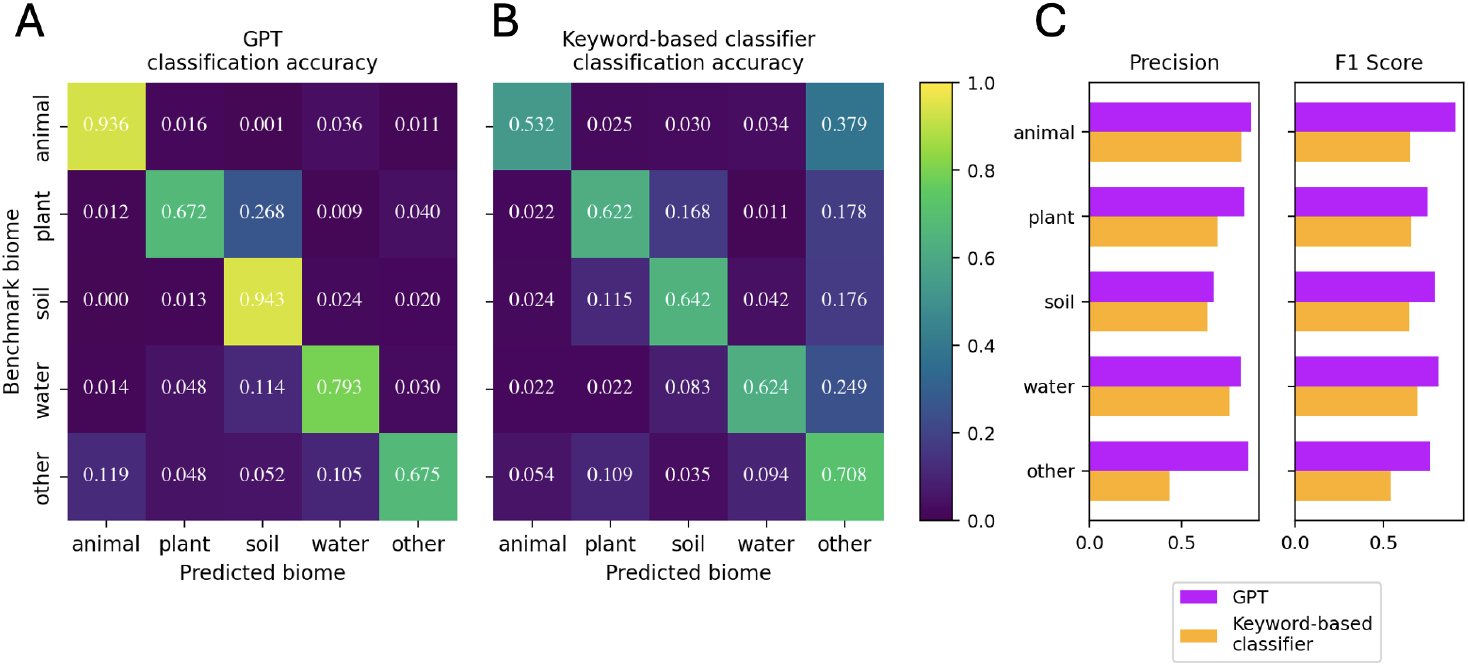
Heatmaps of biome classification accuracy (an average over all GPT runs). Accuracy rates between benchmark and GPT predicted biome classifications and between benchmark and the keyword-based biome classifications are shown in the respective heatmaps. Precision and F1-score bar plots are shown on the right. Notably, the ‘unknown’ category from the previous classification is aligned with ‘other’ from GPT predictions, for direct comparison. The heatmaps are normalized by row. The benchmark dataset consists of *n*=1000 samples. Overall biome accuracy: 80.6% (GPT); 62.5% (Keyword-based classifier). Cohen’s Kappa: 0.757 (GPT); 0.530 (Keyword-based classifier).

Importantly, GPT markedly outperformed MicrobeAtlas’s current, keyword-based classification system (“keyword-based classifier”) in terms of biome prediction accuracy (80.6% vs. 62.5%), precision (81.7% vs. 67.3%) and F1-scores (80.2% vs. 63.5%). We observed the largest performance gain for animal samples (accuracy: 93.6% vs 53.2%, precision: 87.8% vs. 8.24%; F1-score: 90.6% vs. 64.7%), which the keyword-based classifier frequently bins into ‘unknown’. While both methods struggle with plant samples, which they often misclassified as soil (Fig. 2A-B), GPT again produced fewer misclassifications on this subset (4.0% vs. 17.8%). In addition, GPT shows improved predictions in the reverse direction by mislabeling soil samples less frequently as plant biome, compared to the keyword-based classifier (1.3% vs. 11.5%). Overall, GPT predictions resulted in a more balanced distribution across biomes and a better alignment with expected frequencies, as indicated by Cohen’s kappa values (GPT: 0.757; keyword-based classifier: 0.530). (**Figure 2**)

### Analysis of misclassified samples

The overall 19.4% of incorrect predictions led us to investigate whether specific samples are consistently misclassified in predictable ways. Detailed analysis of all GPT misclassifications confirms a subset of samples with significant biases among misclassified biomes (chi-squared: 9364; p-value < 0.001). A large fraction of misclassified ‘animal’ samples (49.9%) are erroneously predicted as ‘water’. In contrast, the majority of misclassifications within ‘plant’ (82.9%) and ‘water’ samples (58.1%) are ‘soil’ (**Supplementary Figure 3**).

The distribution of misclassifications per sample is right-skewed (skewness: 1.32; kurtosis: 0.23), indicating that while most samples have fewer misclassifications, a small subset exhibit disproportionally high values (**Supplementary Figure 4**). Notably, 23 samples (5% of the total) fall at or above the 95th percentile, with 109 or more misclassifications, which is consistent with a right skewed distribution. Across 117 independent classification runs using different parameter settings, the average number of misclassifications per sample is 26.0 (SD: 36.0). Half of the samples are misclassified 5 times or fewer, and three-quarters have no more than 41 misclassifications. The maximum observed was 117, indicating that some samples were misclassified in every run, highlighting extreme cases where GPT predictions consistently disagreed with the curator-assigned biome. (**Supplementary Figure 4**) A detailed examination of such cases includes: a bioreactor digester wastewater sample (SRS994677), a mock community sample (SRS2217033), and two rhizosphere samples (SRS4776621, SRS942824). The bioreactor digester wastewater sample, categorized as ‘other’ by the curator, was incorrectly assigned to ‘water’ by GPT in 109 out of 117 instances. The other eight were correctly assigned as ‘other’. The mock community from a gut microbiome study should have been categorized as ‘other’ due to its laboratory nature, while it was erroneously assigned to ‘animal’ by GPT in 109 out of 112 instances. The other three were correctly assigned. The two rhizosphere samples were misclassified as ‘soil’ by GPT in all but one case.

### Human versus GPT classification accuracy

To evaluate how close GPT’s performance comes to the upper limit of achievable accuracy, we compared it to that of a human annotator. A trained molecular biologist, with no prior exposure to the project, was given the same prompt instructions as GPT and was asked to classify sample biomes and sub-biomes. While against the benchmark dataset, GPT achieved an accuracy of 79.76% (n=499; SD=40.0), the human annotator reached 78.0% (n=250; SD=33.0).

In a second round, both GPT and the human were given the same set of samples to classify, this time using an improved prompt. The only change was an added instruction: “Please note that rhizosphere samples should be categorized as ‘plant’, and sediment samples as ‘water’.” With the improved prompt, both classifiers showed improved performance. GPT’s accuracy increased to 83.17% (SD=37.0), while the human annotator reached 88.0% (SD=33.0).

The improvement in accuracy between GPT’s initial classification and the human’s performance with the improved prompt was statistically significant (adj p-value=0.031), but also between the human’s attempt first attempt (with the initial prompt) and the human’s second attempt (better prompt) (adj p-value≤0.001). No significantly different performances were detected for the sub-biome classification, neither between GPT and the human, nor between prompt versions (adj p-value=1).

### Resource-saving pipeline settings and their impact on output quality

One way to reduce GPT costs is by minimizing the token count submitted to the API, which we achieved by eliminating fields that are empty or have non-informative place-holders such as “NaN” and “unknown”. While the input had to be lengthened in some areas (for example, Environment Ontology *i*.*e*. ENVO codes needed to be converted from numeric IDs to their English text counterparts), we overall achieved a 35% decrease in the size of the input data. Specifically, for the GPT-3.5-turbo-0125 model, which at the time of writing charged $0.5 per million input tokens, a 35% reduction in tokens results in the cost for processing 4 million samples dropping from approximately $1044 to $680.

Instead of sending metadata for individual samples in separate requests, we explored grouping multiple samples’ metadata into single requests (chunking) to reduce the token overhead of repeatedly specifying the input/system prompt. We compared the cost-effectiveness of chunking versus no chunking, and evaluated whether chunking affected performance. We evaluated both biome as well as sub-biome prediction. Our initial findings show no significant performance difference between mild chunking *(i*.*e*.: chunk size 3000: median of 5 samples, maximum of 17 samples per request) and no chunking (one sample per request). This was consistent across both biome prediction (adj *p*-value=0.56) and sub-biome prediction (adj *p*-value=1) (**Figure 3B**)

**Figure 3.**
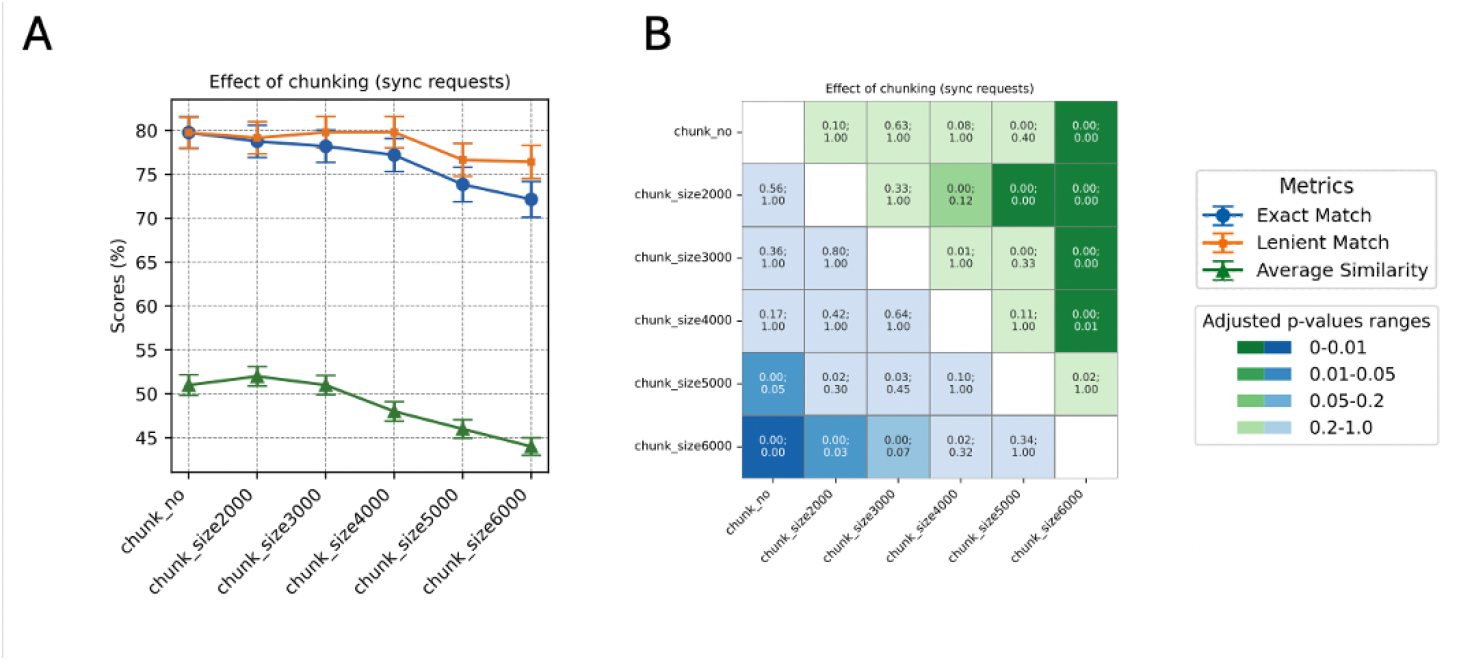
Performance comparison of GPT requests with different chunking sizes. **A**) Accuracy percentages for biome prediction are shown, reflecting both exact matches and lenient matches between the GPT-generated output and the curator-assigned biomes. The similarity for sub-biome prediction is represented through average cosine similarity. Chunk size (2000 up to 6000) refers to the number of tokens within a single request *i*.*e*.: chunk. “Chunk_no” refers to no chunking, where metadata from a single sample is sent as a request. **B**) P-values (top of each cell) and adjusted p-values (bottom of each cell) of the performance comparisons are displayed. Cells shaded in green represent the statistical significance of biome accuracy comparisons, while those in blue denote the significance of sub-biome similarity comparisons. The color intensity varies according to the p-value significance. Mcnemar’s and paired t-tests were performed for biome and sub-biome prediction comparisons, respectively. Bonferroni correction was applied on p-values.

Biome prediction starts to slightly worsen when using larger chunk sizes: biome accuracies for chunk sizes of 2000, 3000, 4000, 5000, and 6000 tokens per chunk (median sample counts of 3, 5, 7, 9, and 15, respectively) were 78.76%, 78.2%, 77.2%, 73.85%, and 72.15% (**Figure 3A**). The noticeable drop in accuracy starting with chunk size 6000 was significant compared to chunk size 2000 (adj p-value=0.03) (**Figure 3B**). Additionally, cosine similarity between benchmark dataset sub-biomes and predicted sub-biomes starts to lower significantly with chunk size 5000 (<quantify>) compared to chunk size 2000 (<quantify>) (adj p-value=0.00115), indicating that GPT is worse at predicting sub-biomes at higher chunk sizes (**Figure 3B**).

### Models & output formats

We compared the performance of three models: GPT-3.5-turbo-0125, GPT-3.5-turbo-1106 and GPT-4-0613 by sending synchronous requests to each model. The accuracy of biome predictions is found consistent across all models, with no statistically significant differences (adj p-value ≥ 0.05) (**Figure 4**). However, we observed differences in the cosine similarity between benchmark dataset sub-biomes and predicted sub-biomes. Specifically, GPT-4-0613 demonstrated a slightly yet significantly higher cosine similarity compared to both GPT-3.5-turbo-0125 (adj p-value ≤ 0.001) and GPT-3.5-turbo-1106 (adj p-value ≤ 0.001) (**Figure 4B**).

**Figure 4.**
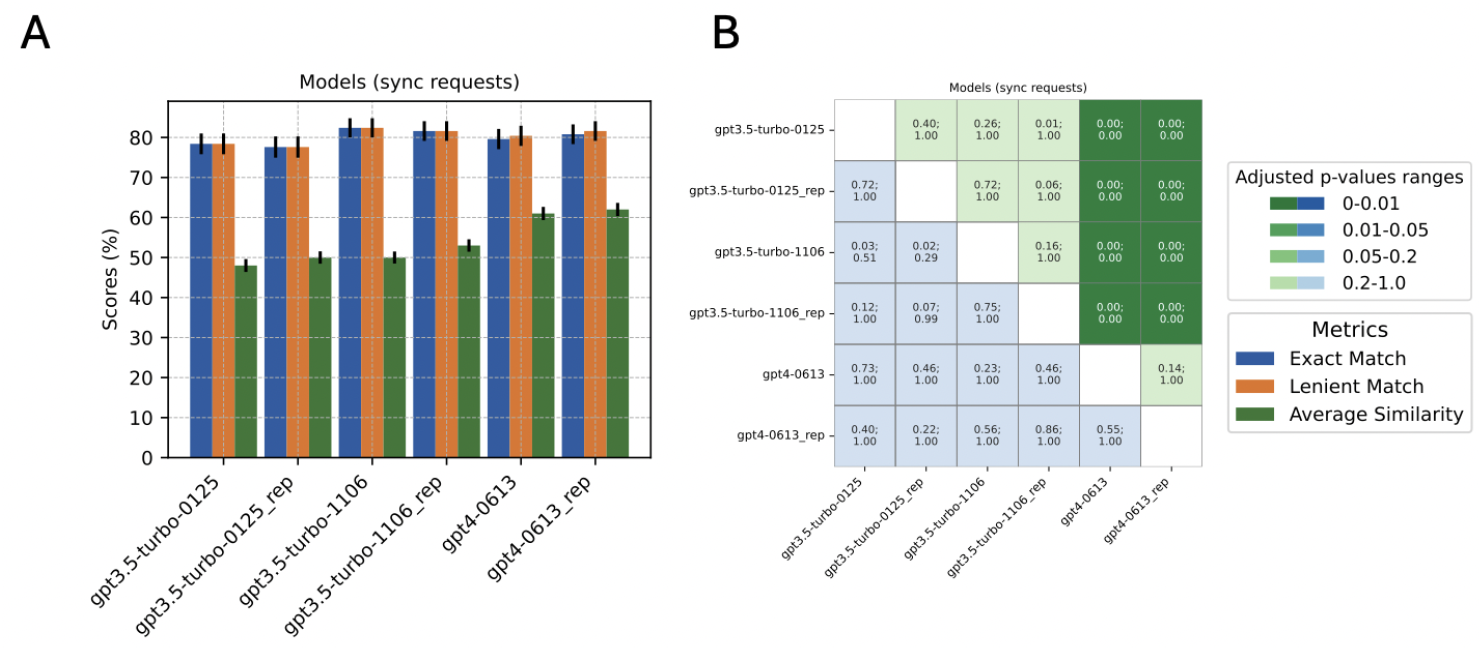
Performance comparison of GPT using different models. **A**) Accuracy scores for biome prediction are shown, reflecting both exact matches and lenient matches between the GPT-generated output and the curator-assigned biomes. The similarity for sub-biome prediction is represented through average cosine similarity. **B**) P-values (top of each cell) and adjusted p-values (bottom of each cell) of the performance comparisons are displayed. Cells shaded in green represent the statistical significance of biome accuracy comparisons, while those in blue denote the significance of sub-biome similarity comparisons. The color intensity varies according to significance. Each run had a replicate (suffix “rep”). Mcnemar’s and paired t-tests were performed for biome and sub-biome prediction comparisons, respectively. Bonferroni correction was applied on p-values.

To assess the impact of different output formatting on GPT’s performance, we designed two system prompts that instruct GPT to format its output either inline (using double underscores to separate answers for each sample), or in json format (by stating it in the prompt and by specifying the ‘format’ parameter in the API request). While biome prediction accuracy remained constant across both formats, we observed an improvement in sub-biome cosine similarity when responses were formatted in json (adj p-value < 0.001). (**Figure 5**)

**Figure 5.**
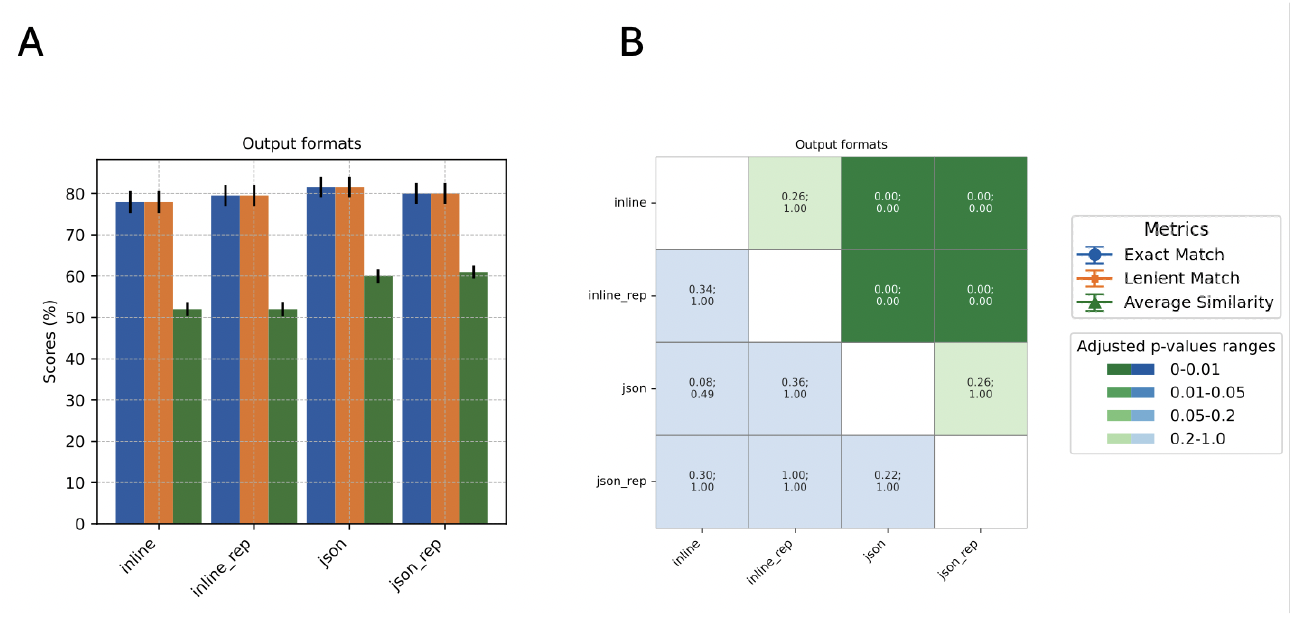
Performance comparison of GPT using either in-line or json output format. A) Accuracy scores for biome prediction are shown, reflecting both exact matches and lenient matches between the GPT-generated output and the curator-assigned biomes. The similarity for sub-biome prediction is represented through average cosine similarity. B) P-values (top of each cell) and adjusted p-values (bottom of each cell) of the performance comparisons are displayed. Cells shaded in green represent the statistical significance of biome accuracy comparisons, while those in blue denote the significance of sub-biome similarity comparisons. The color intensity varies according to the p-value significance. Mcnemar’s and paired t-tests were performed for biome and sub-biome prediction comparisons, respectively. Bonferroni correction was applied on p-values.

### The effect of tweaking creativity parameters

Parameters such as temperature, nucleus sampling, and penalty settings influence how deterministic or diverse the outputs of LLMs are, and thus could potentially affect prediction. Adjustments to temperature, nucleus sampling, or presence-penalty settings did not impact the prediction accuracy for biome or sub-biome categories (adj p-value ≥ 0.05) (**Supplementary Figure 5**). Accuracy consistently dropped with increasing frequency penalty, from an initial 79.2% (freqp 0.0) to a minimum of 64.3% (freqp 2.0). This decline was significant (p-adj < 0.001) (**Figure 6**). However, when the more lenient criterion for string-matching of the GPT output with the benchmark dataset biome was applied (*i*.*e*.: “lenient match”), the impact of increased frequency penalty on biome accuracy was mitigated (freqp 0.0: 80.76%; freqp 2.0: 80.96%) (**Figure 6A**). By examining GPT answers, it was clear that a higher frequency penalty led GPT to deviate from strictly adhering to the prompt instructions “Answer with exactly one word from the following…”. Instead, the model tended to provide unsolicited information *e*.*g*.: instead of providing “animal” for an answer, it provided “animal (incl. human)”. In fact, biome accuracy under the lenient matching condition remained unaltered with differing frequency penalties.

**Figure 6.**
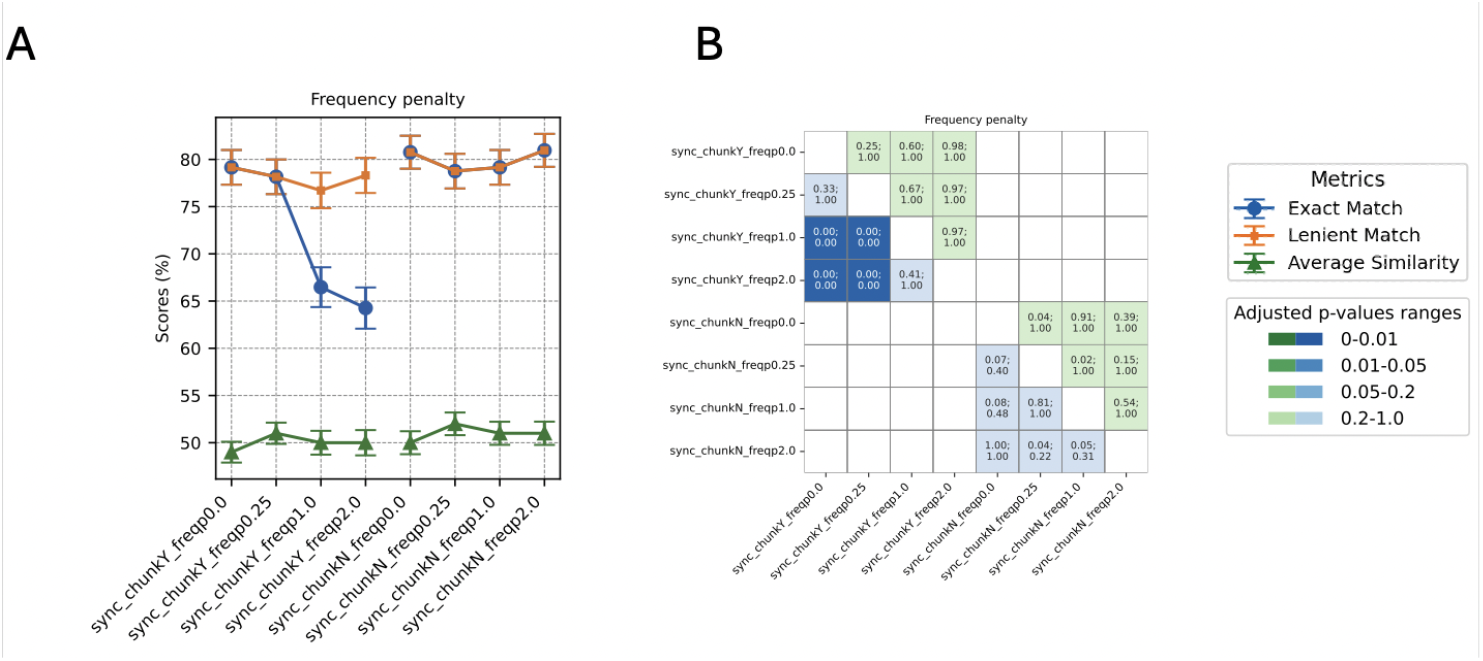
Performance comparison of GPT when tweaking frequency penalty. **A**) Accuracy scores for biome prediction are shown, reflecting both exact matches and lenient matches between the GPT-generated output and the curator-assigned biomes. The similarity for sub-biome prediction is represented through average cosine similarity. **B**) P-values (top of each cell) and adjusted p-values (bottom of each cell) of the performance comparisons are displayed. Cells shaded in green represent the statistical significance of biome accuracy comparisons, while those in blue denote the significance of sub-biome similarity comparisons. The color intensity varies according to the p-value significance. For an assessment of all other creativity parameters see Supplementary Figure 3. Mcnemar’s and paired t-tests were performed for biome and sub-biome prediction comparisons, respectively. Bonferroni correction was applied on p-values.

Remarkably, significant differences across creativity parameters were detected only in the case of chunked requests. Without chunking, neither synchronous requests nor asynchronous requests showed significant differences (**Supplementary Figure 6**). This indicates that the tweaking of creativity parameters, specifically frequency penalty, affects the output only when multiple samples’ metadata are submitted in a single request.

### Synchronous and asynchronous requests

To assess the reliability of different querying strategies in large-scale automated analyses, we compared the robustness of synchronous (without chunking) and asynchronous requests. We conducted 10 replicate tests for each: 250 randomly selected metadata samples were sent individually to 10 synchronous runs or 10 asynchronous runs (replicates 1 to 10). For synchronous requests, the biome accuracy (lenient match) did not vary (range=0.79-0.81), and neither did the sub-biome cosine similarity (range=0.49-0.50) (**Supplementary Figure 6A**). Asynchronous requests demonstrated similar consistency for biomes accuracy (range=0.82-0.83), and no significant variations across different runs for sub-biome predictions (range=0.56-0.58) (adj p-value ≥ 0.05) (**Supplementary Figure 6B**).

We evaluated the performance differences between synchronous and asynchronous requests in terms of both biome and sub-biome prediction accuracies. Our analysis revealed that whether requests are sent synchronously or asynchronously does not correlate with a better prediction of biomes nor sub-biomes (adj p-value=1) (**Supplementary Figure 6C,F**).

### Geographic location prediction performance

The various GPT models we tested (*i*.*e*.: GPT-3.5-turbo-0125, GPT-3.5-turbo-1106, and GPT-4-0613) were also instructed to extract the geographical location where each sample was collected, including the country (in textual format, *i*.*e*. not in latitude/longitude coordinates). We measure the performance by comparing this output with textual descriptions derived directly from lat/lon coordinates parsed from the metadata. Among 130,689 samples analyzed, 95.94% matched the geo-coded location. The distribution of the matching and mismatching samples can be visualized on a global map (**Figure 7**).

**Figure 7.**
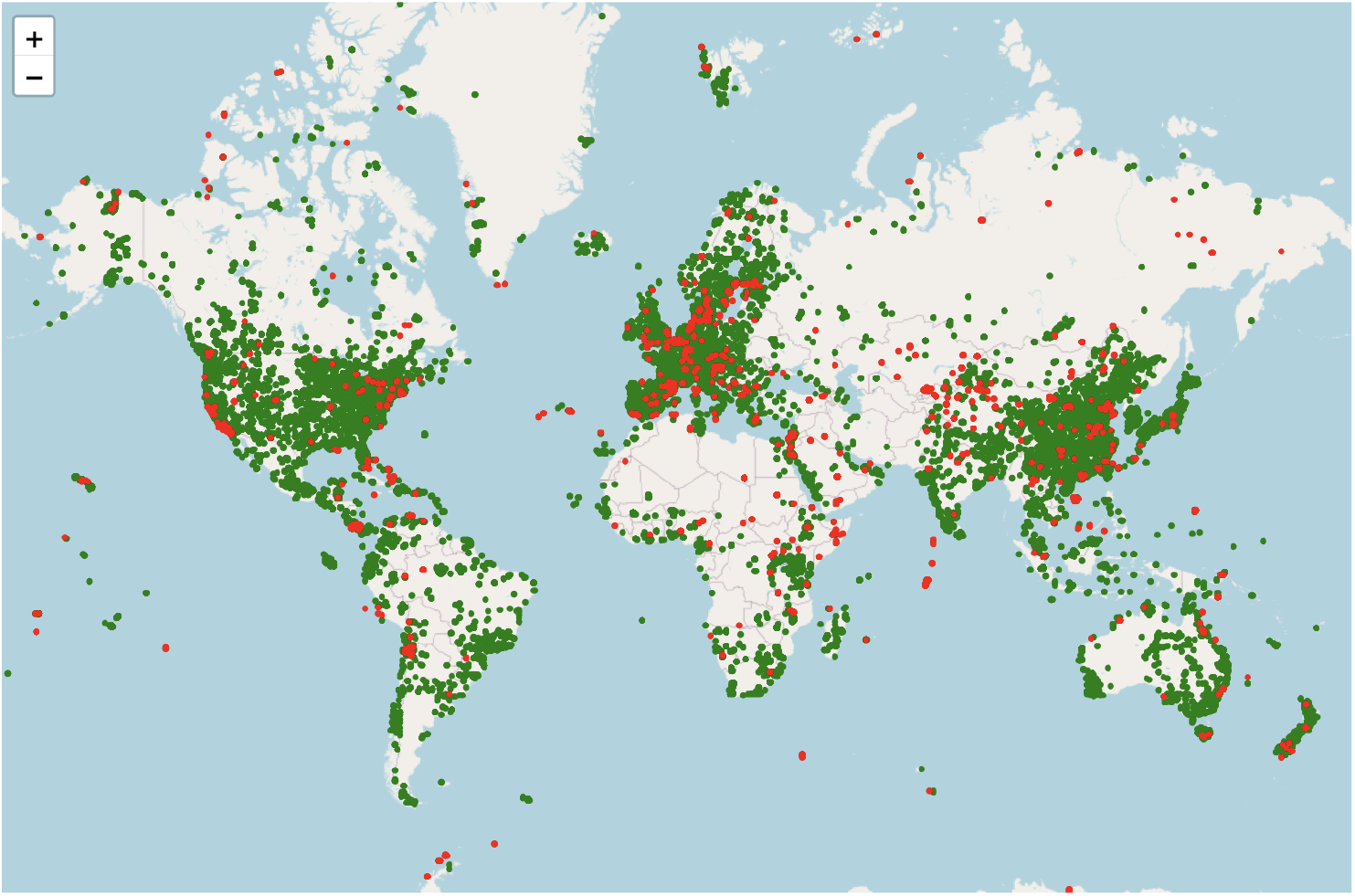
Distribution of samples with matching (green) and mismatching (red) locations (GPT-predictions *vs*. metadata-extracted geo locations). Link to full image: 10.5281/zenodo.15274397

To further analyze the 4.06% mismatches (n=5,312), we obtained the actual coordinates for the GPT-predicted locations using the GoogleMaps API and assessed the distance between these and the metadata-derived coordinates. The distances ranged from 0.004 km to 19,856 km (1^st^ quartile: 129.03 km, 3^rd^ quartile: 9,900.94 km). The distribution of the mismatching samples color-coded by distance categories can be visualized on a global map (**Supplementary Figure 7**). For a focused investigation, a subset of 100 misclassified samples, randomly picked and with a discrepancy above 1000 km between the GPT-predicted locations and the metadata-derived coordinates, were examined manually. This analysis revealed that :

- In half of the cases GPT correctly predicted the location, while the lat/lon coordinates parsed from the metadata were incorrect. This happened for various reasons: the coordinates in the metadata were either completely wrong (30%), lat/lon were swapped (6%), had wrong longitudinal signs (negative (34%) or positive (6%)), had wrong latitudinal signs (negative, 10%), a wrong latitude (6%), or the coordinates pointed at the academic institution of the authors rather than to the location where the sample was collected (8%);
- In 27% of the cases both GPT and coordinates pointed at the right geographic location, but the name given by GPT did not textually match with the geographic location from the coordinates (*e*.*g*.: McMurdo station vs Antarctica). This, for example, often happened with coastal samples of the United States, where GPT returns “United States”.
- In 16% of the cases the coordinates pointed to the right location, but GPT didn’t extract it correctly. This happened because GPT mistook the institute location for the sampling location (37.5%), the coordinates pointed to a water body or an island (*e*.*g*.: “Hawaii)” while GPT predicted the location after the country of belonging of that water body (*e*.*g*.: “United States”) (18.8%), or the metadata did not clearly mention the location in text format (43.8%).;
- In 7% of the cases metadata-extracted coordinates and GPT geo location did not match because neither information was retrievable from the metadata as it was ambiguous or absent.

### Informative metadata fields

Not all metadata fields contain information about sample origins, but as field names are not always standardized, it can be difficult to know *a priori* which fields to parse and focus on. By programmatically comparing the metadata of 1000 samples to their curator-assigned sub-biomes, we determined which fields typically report the sample origin. We found that the sample origin was documented in 67 distinct fields when considering full matches (*e*.*g*.: “cow rumen”) and in 127 distinct fields when considering partial matches (*e*.*g*.: “cow” or “rumen”). On average, the full origin of a sample is found in 1.36 fields per sample (SD: 1.67), and at least part of the sample origin is reported in 3.0 fields (SD: 2.2) per sample, indicating a modest variability in how sample origins are documented across different metadata fields.

Some fields are more commonly used than others to report the origin of the sample. The fields most frequently containing sample origin information are: “study_STUDY_ABSTRACT” (267 instances), “sample_SCIENTIFIC_NAME” (225 instances), “study_STUDY_TITLE” (212 instances), “sample_isolation_source” (113 instances), and “study_STUDY_DESCRIPTION” (53 instances). Despite their relevance for mentioning sample origins, the fields “study_STUDY_ABSTRACT” and “study_STUDY_DESCRIPTION” are more challenging to parse given their average word count (72.8 and 144.8, respectively) compared to “sample_SCIENTIFIC_NAME” (2.12), “sample_isolation_source” (2.9), and “study_STUDY_TITLE” (8.9).

The variability in field usage does not seem to be arbitrary, but appears influenced by the biome associated with the sample. For example, sample origin is more likely to be found under the field “sample_host” in the case of animal and plant samples, while the fields “sample_env_biome” and “sample_env_feature” are more likely to contain useful information in the case of soil, water and “other” samples. This demonstrates that certain fields are preferentially selected to report sample origin, depending on the sample biome. (**Supplementary Figure 8**)

## Discussion

A central goal of data repositories is to enable the effective secondary use of research data, potentially leading to new discoveries beyond the original scope of the study. In practice, however, most datasets are rarely re-analysed by researchers outside the submitting lab. A major barrier to reusability is poor metadata–often incomplete, inconsistent, or non-standardised–which limits automated parsing and interpretation. As a result, the full potential of many datasets remains untapped. Two strategies come to mind to address this issue: (1) enforcing more rigorous reporting standards at the time of submission, and (2) retroactively improving metadata quality using computational tools. While ongoing efforts have improved metadata submission guidelines, these changes are difficult to enforce consistently and do not address the vast amount of legacy data already in repositories. In contrast, large language models (LLMs) offer a flexible and scalable means to retrospectively annotate and standardize metadata, even when it is noisy or poorly structured. In this study we evaluate the use of one such model–GPT–for metadata re-annotation of environmental sequencing samples.

When evaluating GPT’s performance in classifying environmental sequencing samples, we found that it consistently outperformed a traditional keyword-based classifier. Unlike the latter, which relies on static keyword matching and extensive hard-coded white lists, GPT leverages context and semantics to disambiguate terms based on their usage and surrounding information within the metadata. To better contextualize GPT performance, we compared it to that of a trained molecular biologist with no prior exposure to the project. Both were presented with the same metadata and prompt, and their classifications were evaluated against the benchmark. Under standard conditions, GPT and the human annotator achieved similar accuracy. When given a clearer, more explicit prompt, both improved. This demonstrates not only GPT’s capacity to perform at a near-expert level for this classification task, but also highlights the importance of precise instructions—an aspect that benefits both human and machine annotators.

However, GPT is not without its limitations. One challenge is the need to translate ontology accession codes—commonly found in metadata—into human-readable terms, as LLMs do not interpret these codes natively. Secondly, LLMs do not always “read between the lines”. For instance, if a sample is part of an animal study but correctly described as a mock community, GPT may still classify it as ‘animal’ rather than ‘other’. Prompt adherence is also imperfect: if the metadata points frequently at the term ‘soil’, GPT may default to assigning that biome, regardless of more nuanced cues. This was especially evident for rhizosphere and sediment samples, which were major contributors to biome misclassification. These cases illustrate a broader challenge: sample classification in environmental data is often inherently ambiguous. A sediment sample taken from a coastal area, for example, could reasonably belong to either ‘soil’ or ‘water’ depending on phrasing. Similar issues affected the keyword-based classifier, which also showed bias in such edge cases. These examples underscore the difficulty of resolving ambiguity in free-text metadata, regardless of the method applied.

Taken together, the ∼20% misclassification rate observed with GPT likely reflects an upper limit LLM performance, rather than a true failure rate. Given the unavoidable ambiguity and challenges in re-annotating diverse samples against a necessarily limited and simplified classification scheme, even a second human curator might not agree with the benchmark labels. This suggests that even LLMs operating at human level may not be sufficient for perfect categorization–especially in domains like environmental sampling where a certain amount of overlap is to be expected.

An alternative to direct classification is to prompt the model to generate a brief description of the sample’s origin and then derive embeddings from these summaries. These embeddings can be compared to those generated from curator-provided descriptions. This approach allows for more flexible sample grouping: even if a sample is misclassified at the biome level, its sub-biome description—*e*.*g*., “mock community”—may still position it correctly in embedding space alongside similar samples. In this study, we used such comparisons primarily to validate GPT’s output via cosine similarity scores, but the same method could also support clustering or reclassification of ambiguous samples.

Tweaking LLM ‘creativity parameters’ such as temperature, nucleus sampling, and presence penalty had no impact on biome or sub-biome prediction accuracy. However, increasing the frequency penalty led to reduced biome accuracy—but only when multiple samples were included in a single request (*i*.*e*., chunking). This effect disappeared when samples were submitted individually. A likely explanation is that repeated use of a term like ‘animal’ within a chunk triggers the penalty, prompting GPT to vary its wording (*e*.*g*., returning “animal (human)” instead). Supporting this, the accuracy drop vanished when using lenient string matching. Interestingly, rather than avoiding penalized terms altogether, GPT tended to modify them. Aside from frequency penalty effects during chunking, no other creativity parameter significantly affected classification performance.

We also assessed the impact of request mode, model version, and output format on GPT’s performance. As expected, there was no difference between synchronous and asynchronous requests, since both use the same underlying model. Given this, asynchronous requests are currently preferable, as they are easier to manage at scale and are more cost-effective. Model choice, however, did influence output: GPT-4 outperformed both GPT-3.5-turbo-1106 and GPT-3.5-turbo-0125 in sub-biome prediction. Output format also had an effect. While biome predictions were unchanged, sub-biome accuracy improved significantly when using JSON rather than inline formatting. Although both formats yielded similar recall, the structured format may help the model better focus on the task, possibly by encouraging more consistent output or reducing ambiguity during generation.

As the use of GPT’s API is invoiced based on the number of input and output tokens, we attempted to reduce costs by (1) cleaning away uninformative fields from the metadata and (2) sending multiple samples’ metadata under the same request (*i*.*e*.: chunking). The preliminary cleaning of the metadata proved valuable, reducing input token size by 35%. We hypothesized that sending too many samples at once might “confuse” the model and tested the limit of this strategy. In our case performance declined significantly at 5000 tokens per request (roughly nine samples), while remaining stable at 2000 tokens (about three samples). A decline was already visible at 4000 tokens (seven samples), although it did not reach statistical significance. Although we did not test chunking with asynchronous requests, we have no reason to believe the results would differ. It is important to note that these thresholds may vary for different metadata types.

Our analysis of 1000 samples demonstrated that the sample origin typically appears in 1 to 3 fields but spans across no fewer than 127 distinct field types, underscoring the immense variability in how metadata is reported. The best field for inferring the broad origin of a sample is the study_STUDY_ABSTRACT field, which is also the most verbose and thus challenging to process. The sample_SCIENTIFIC_NAME field often reflects the sample origin in host-associated metagenomic samples and is more concise in length. Overall, the usefulness of specific fields varies across biomes, suggesting that a one-size-fits-all selection of metadata fields would not perform equally well across all sample types. This highlights the persistent challenges in standardizing metadata extraction, despite ongoing efforts to regulate SRA submissions.

We also used GPT to extract the geographic location of each sample’s collection site. This was useful in cases where latitude/longitude coordinates were missing, incorrect, or inconsistently formatted. Issues included swapped coordinates, decimal errors, or identical shifts across samples of the same study due to submitters using the drag-copy function in Excel. In many such cases, GPT successfully inferred the sampling country or locality based on free-text metadata, achieving close to 96% accuracy. Among mismatches, about half were due to incorrect metadata coordinates, while a quarter were semantic mismatches (*e*.*g*., “Antarctica” vs. “McMurdo Station”). In a fifth of the cases, GPT misidentified the location, either due to confusing institution names with sampling sites or due to vague metadata. A small fraction of samples had no retrievable geographic information at all.

Overall, our results indicate that the context-aware parsing capabilities of LLMs are sufficient for metadata (re-)annotation—at least in relatively structured tasks like microbiome sample origin classification. Errors still occur, partly due to model limitations, partly due to the inherent ambiguity of real-world samples, and occasionally due to parsing issues. More straightforward technical constraints include rate limits imposed by commercial APIs and token limits per request. These will vary depending on the model used, and we did not test free or open-access models—an acknowledged limitation of our study. Nevertheless, we suggest that LLMs like GPT can usefully complement existing workflows across the data management lifecycle, including guiding metadata submission or assisting with post-hoc metadata curation. Good quality metadata is essential for enhancing data reusability, will be crucial in managing the growing volume of microbiome data, and will ultimately support new discoveries in the field.

## Data availability

Source code is available on Github (metadata_mining/tree/main/scripts). Prompts and benchmark dataset are available on Github (metadata_mining/tree/main/source_data). All results of biome and subbiome validation and all results from the comparisons performed are available on Github (biome_subbiome_results.csv, and biome_subbiome_stats.csv, respectively). GPT answers used to evaluate accuracy (10.5281/zenodo.15274823) and interactive plots (10.5281/zenodo.15274397; 10.5281/zenodo.15274467) are available on Zenodo.

## Acknowledgements

We are grateful to the members of the von Mering lab—past and present—for their valuable input and stimulating discussions that shaped the development of this manuscript, in particular Maria Dmitrijeva, João Rodrigues, Tao Fang, Tülay Karakulak, Damian Szklarczyk, Qingyao Huang, Matteo Peluso, Radja Hachilif, and Maria Heimlicher. We extend special thanks to George Hausmann for his critical role in annotating a subset of benchmark samples. His independent annotations provided an important reference point for evaluating the upper bound of classification performance and greatly contributed to the interpretation of results. D.G., D.P. and C.v.M. were supported by the Swiss National Science Foundation through its National Competence Center for Research “Microbiomes” (grant number 310030_192567). L.M., N.N., and E.P. were supported by an SNSF project grant to C.v.M. (grant 310030_192569).

## Supplementary Figures

**Supplementary Figure 1:**
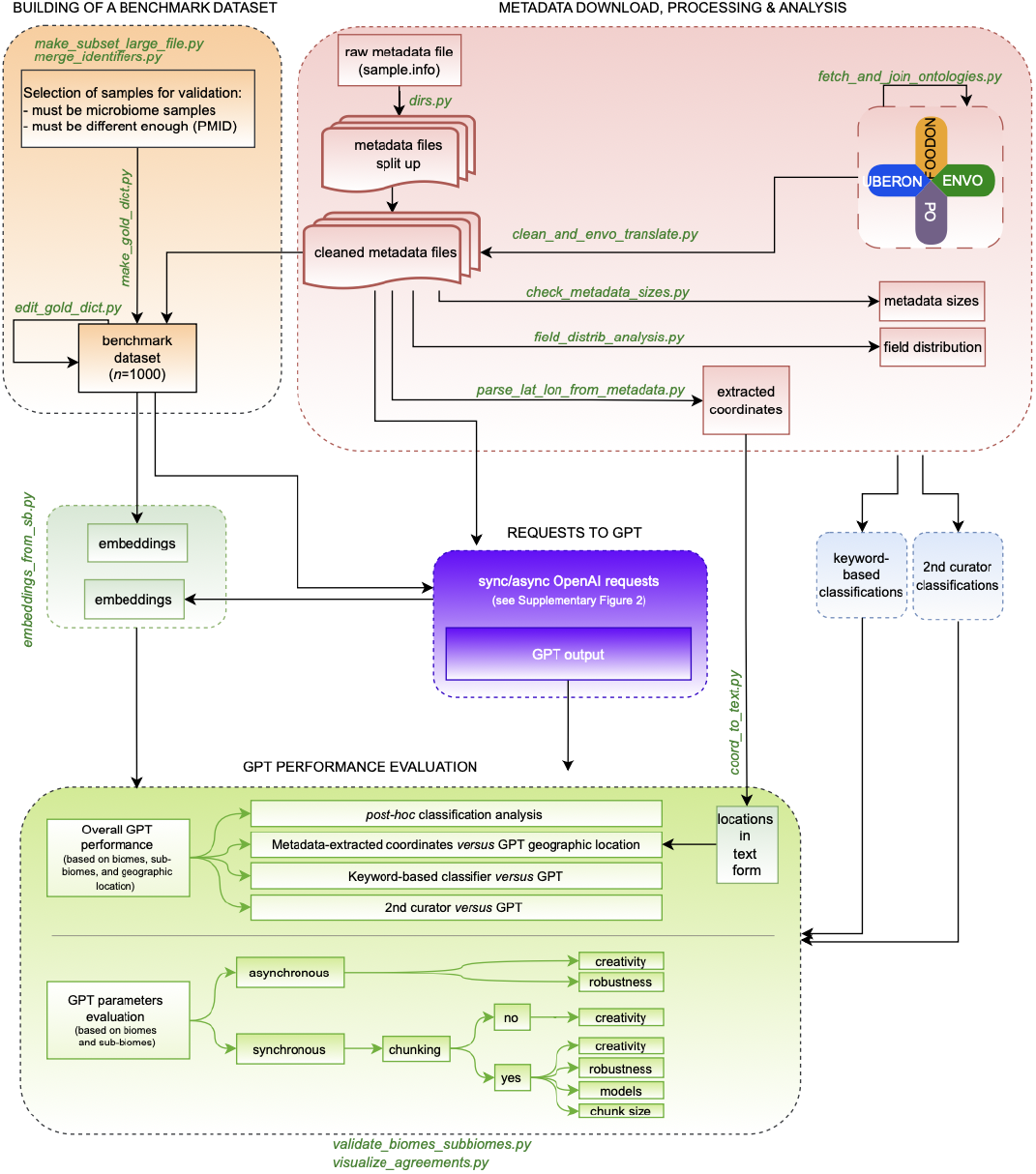
Pipeline in detail.

**Supplementary Figure 2.**
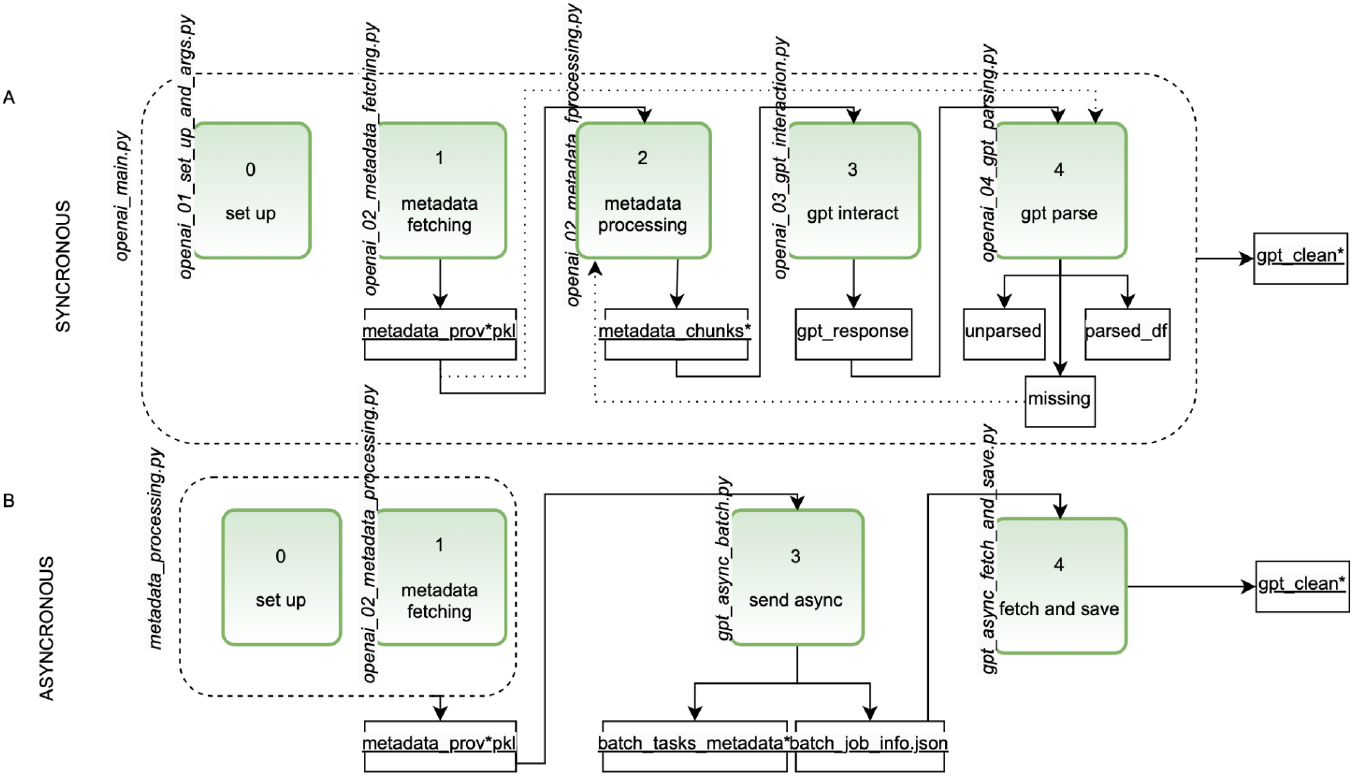
Pipeline of synchronous (**A**) and asynchronous (**B**) requests.

**Supplementary Figure 3.**
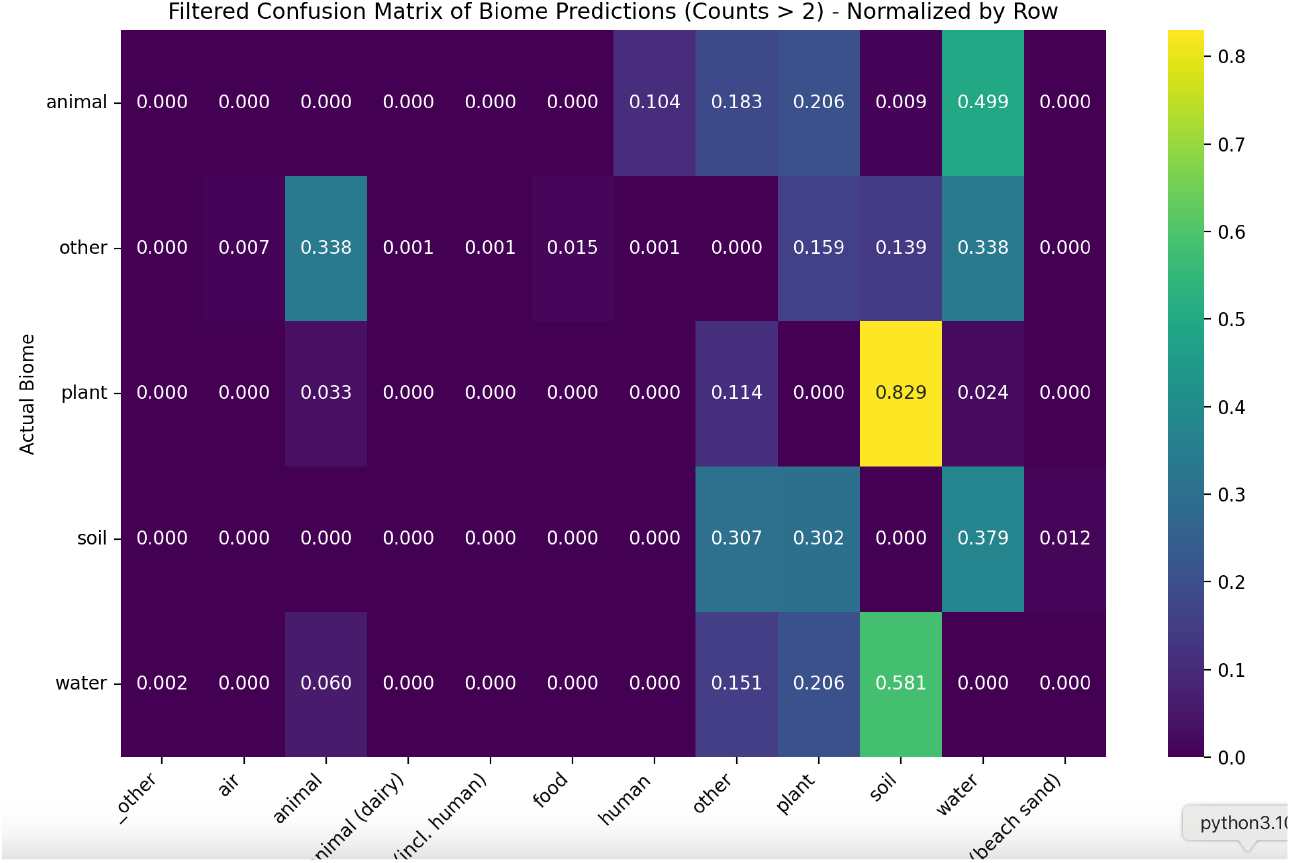
Misclassifications.

**Supplementary Figure 4.**
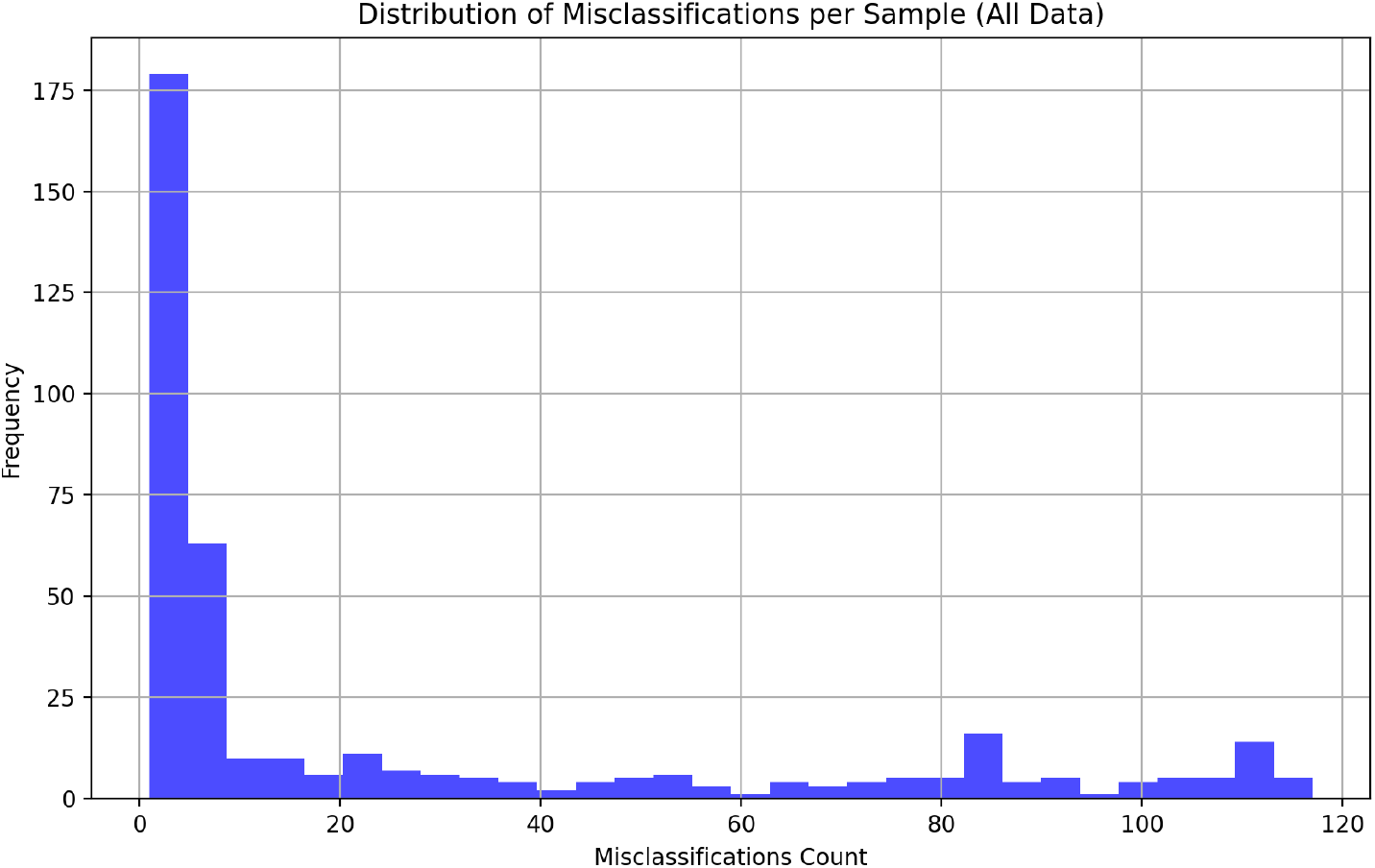
Distribution of sample misclassifications by GPT.

**Supplementary Figure 5.**
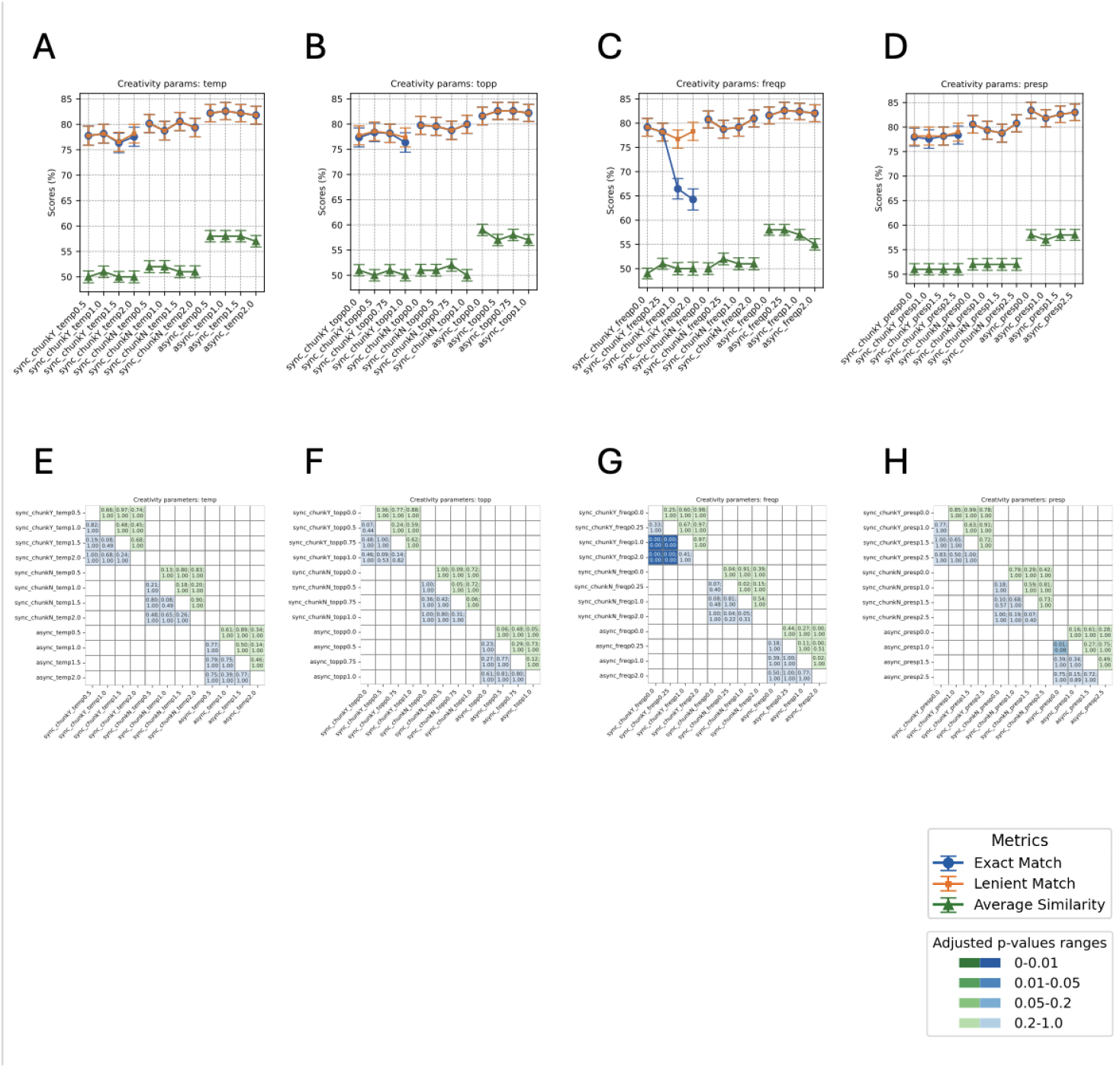
Performance comparison of GPT when tweaking creativity parameters: temperature (temp), nucleus sampling (topp), frequency penalty (freqp) and presence penalty (presp). **A-D**) Accuracy scores for biome prediction are shown, reflecting both exact matches and lenient matches between the GPT-generated output and the curator-assigned biomes. The similarity for sub-biome prediction is represented through average cosine similarity. **E-H**) P-values (top of each cell) and adjusted p-values (bottom of each cell) of the performance comparisons are displayed. Cells shaded in green represent the statistical significance of biome accuracy comparisons, while those in blue denote the significance of sub-biome similarity comparisons. The color intensity varies according to the p-value significance. Mcnemar’s and paired t-tests were performed for biome and sub-biome prediction comparisons, respectively. Bonferroni correction was applied on p-values.

**Supplementary Figure 6.**
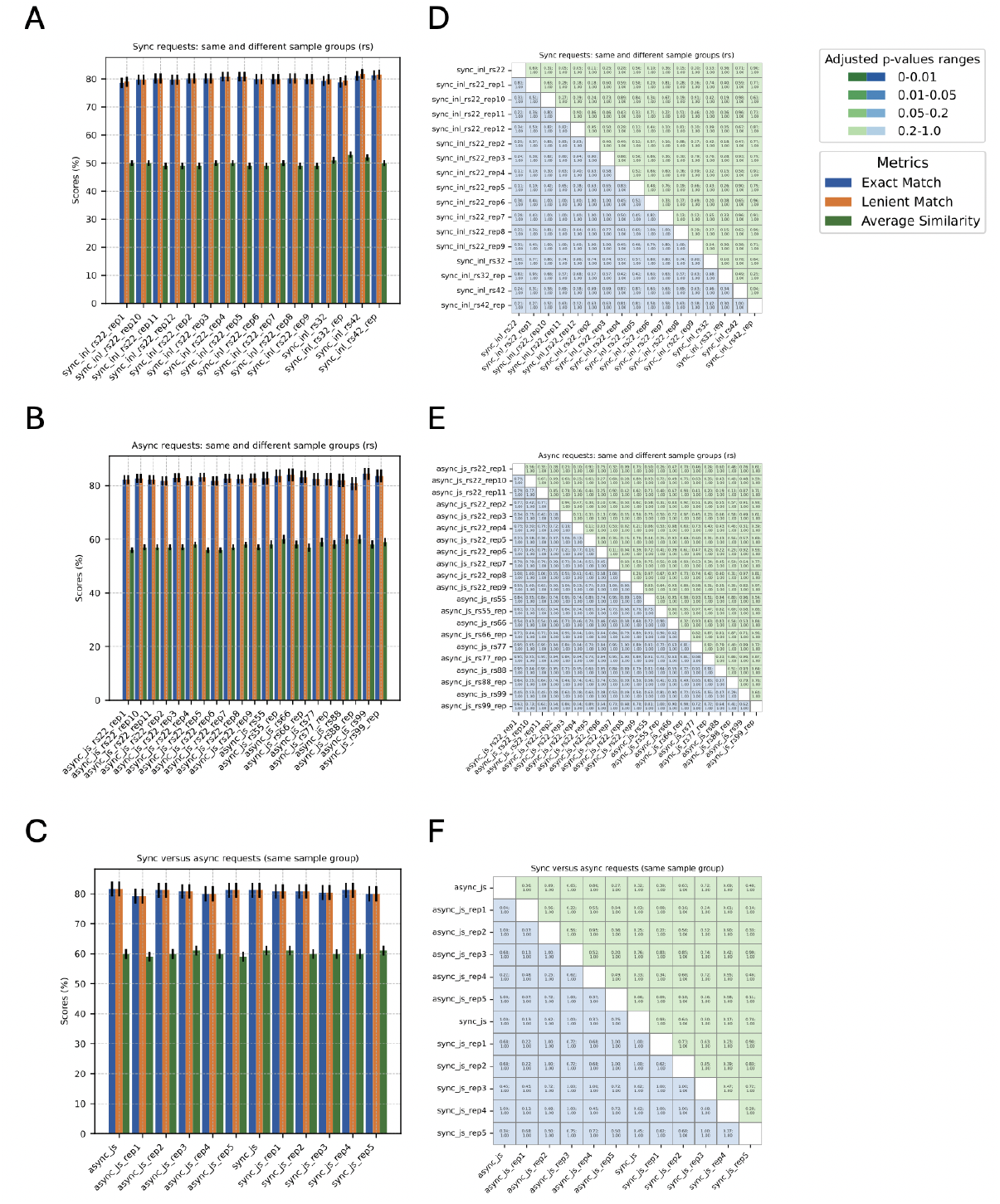
Performance reproducibility. **A-C**) Accuracy scores for biome prediction are shown, reflecting both exact matches and lenient matches between the GPT-generated output and the curator-assigned biomes. The similarity for sub-biome prediction is represented through average cosine similarity. **D-F**) P-values (top of each cell) and adjusted p-values (bottom of each cell) of the performance comparisons are displayed. Cells shaded in green represent the statistical significance of biome accuracy comparisons, while those in blue denote the significance of sub-biome similarity comparisons. The color intensity varies according to the p-value significance. Mcnemar’s and paired t-tests were performed for biome and sub-biome prediction comparisons, respectively, when the sample groups being compared were the same (same ‘rs’ *i*.*e*.: random seed). When comparing different sample groups, independent t-tests were applied. Bonferroni correction was applied on p-values.

**Supplementary Figure 7.**
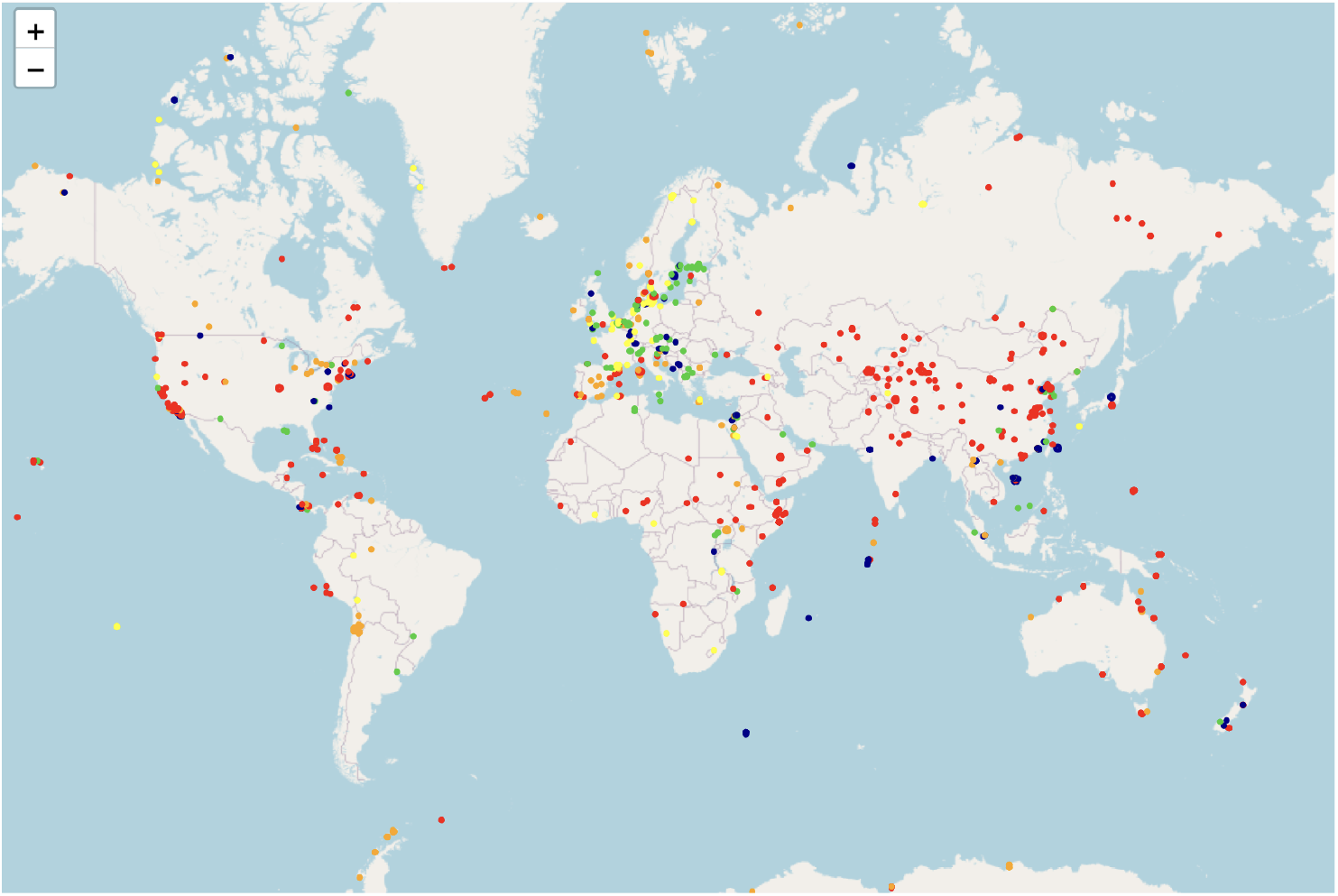
Distribution of the mismatching samples color coded by distance between GPT predicted location and coordinates-extracted geo location (less or equal to 100km: light blue; between 100 and 500 km: green; between 500 and 1000 km: yellow; between 1000 and 4000 km: orange; more than 4000 km: red). Link to interactive figure: 10.5281/zenodo.15274467

**Supplementary Figure 8.**
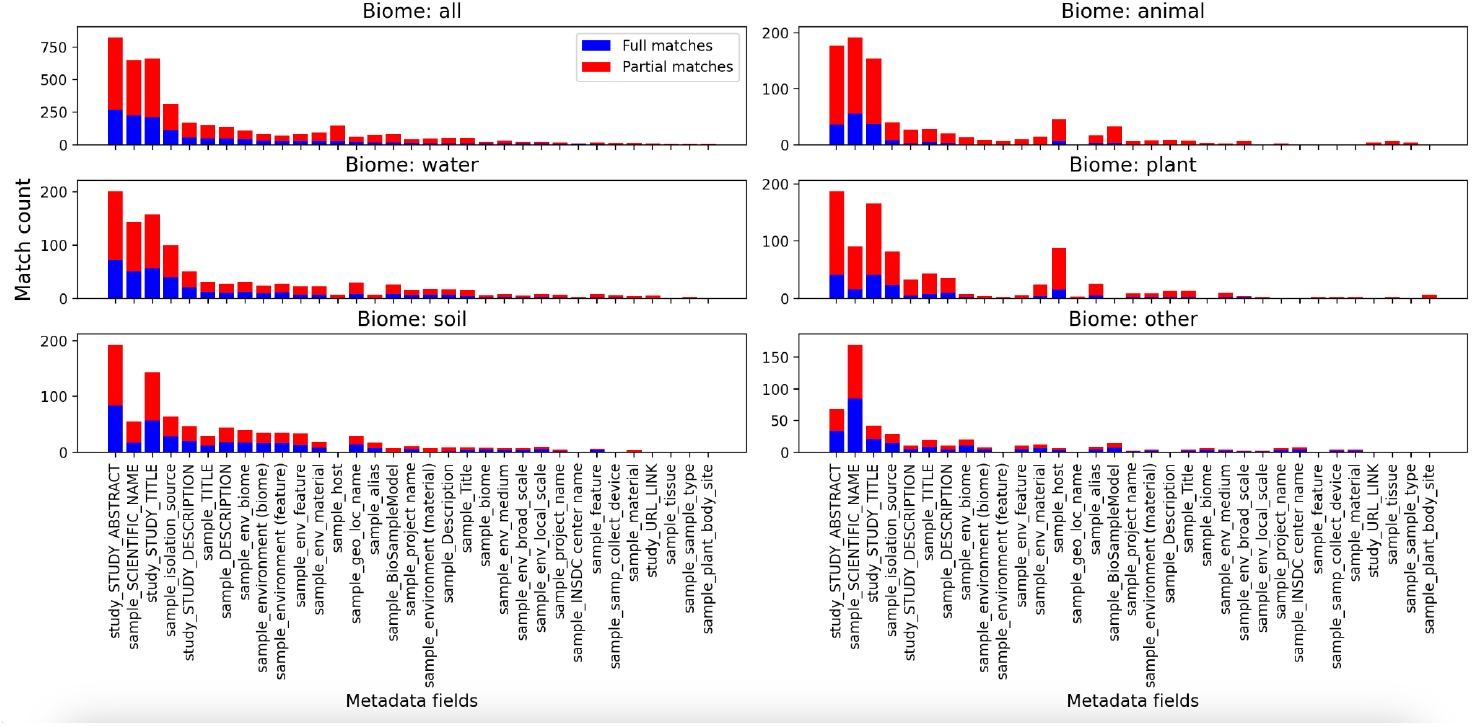
Metadata field distribution. Frequency of metadata fields matching benchmark sub-biomes by biome category. Fields with less than 20 partial matches are filtered out.

